# First record of ‘plasticrusts’ and ‘pyroplastic’ from the European Mediterranean Sea

**DOI:** 10.1101/837484

**Authors:** Sonja M. Ehlers, Julius A. Ellrich

**Affiliations:** Department of Animal Ecology, Federal Institute of Hydrology, Am Mainzer Tor 1, 56068 Koblenz, Germany; Institute for Integrated Natural Sciences, University of Koblenz-Landau, Universitätsstraße 1, 56070 Koblenz, Germany; Independent Researcher, Sankt-Josef-Straße 25, 56068 Koblenz, Germany

**Keywords:** Plastic debris, Rocky intertidal, Sandy beach, Giglio island, Tyrrhenian Sea, Italy

## Abstract

We report the presence of ‘plasticrusts’ and ‘pyroplastic’ from coastal habitats in Giglio island, Tyrrhenian Sea, Italy. These novel plastic debris types have only recently been described for the first time from Madeira island (NE Atlantic Ocean) and the United Kingdom, respectively. While ‘plasticrusts’ are generated by sea waves smashing plastic debris against intertidal rocks, ‘pyroplastic’ derives from (un)deliberately burnt plastic waste. Using Fourier-transform infrared (FTIR) spectroscopy, we identified the ‘plasticrust’ material as polyethylene (PE) and the ‘pyroplastic’ material as polyethylene terephthalate (PET). These polymers are widely used in everyday products and, therefore, contribute heavily to plastic pollution in aquatic and terrestrial environments worldwide. Furthermore, our field surveys suggest that ‘plasticrust’ abundance is related to wave-exposure and that the ‘pyroplastic’ derived from beverage bottles which we frequently found along the Giglio coast. Overall, our findings corroborate the notion that ‘plasticrusts’ and ‘pyroplastic’ are common debris types in marine coastal habitats.

## Introduction

Plastic materials have become part of everyday life and are appreciated for their high durability in single-use items such as beverage bottles, food packaging and shopping bags. However, mismanaged plastic debris is rapidly accumulating in marine (Andrady 2011), freshwater (Wang et al. 2017, Akindele et al. 2019) and terrestrial ecosystems (Scheurer and Bigalke 2018, Panebianco et al. 2019) and can even be found in the most remote locations such as the deep sea (Woodall et al. 2014) and the arctic (Bergmann et al. 2019, Kanhai et al. 2019). Due to the omnipresence and persistence of plastic materials, the term 'Plasticene' has been introduced (Reed 2015). In marine ecosystems, plastic debris has been detected since the 1970s (Carpenter et al. 1972, Carpenter and Smith 1972). To date, most studies on marine plastic pollution examined pelagic environments (e.g. Chambault et al. 2018, Campanale et al. 2019). In contrast, there has been surprisingly little research on plastic debris in rocky intertidal habitats. This is remarkable since marine plastic debris is regularly washed ashore and can easily be fragmented when crashed against intertidal rocks by sea waves (Gestoso et al. 2019). Recently, plastic debris encrusting the Madeira island (NE Atlantic Ocean) rocky intertidal has been termed ‘plasticrusts’ and described as a new type of marine plastic debris that may harm invertebrates grazing on intertidal rocks. Moreover, observations suggest that ‘plasticrusts’ can persist over years and that ‘plasticrust’ abundance may increase over time (Gestoso et al. 2019). Another novel type of plastic debris is ‘pyroplastic’ which has been reported from beaches in the SW United Kingdom recently (Turner et al. 2019). ‘Pyroplastics’ have a stone-like appearance and contain encapsulated plastic debris. Such ‘Pyroplastics’ are formed when plastic waste is burnt and cooled down which creates grayish agglomerates with encapsulated plastic material that may leach toxic substances. Due to their appearance, ‘Pyroplastics’ are difficult to distinguish from natural rocks (Turner et al. 2019). However, to consider novel plastic debris types in marine conservation policy (Gestoso et al. 2019) it is necessary to investigate whether ‘plasticrusts’ and ‘Pyroplastics’ occur on other coasts as well. Since plastic pollution in the Tyrrhenian Sea (European Mediterranean) is relatively high (Campanale et al. 2019), we examined for the first time the occurrence of ‘plasticrusts’ and ‘Pyroplastics’ on Tyrrhenian Sea coasts.

## Materials and Methods

### Studied coasts

On 19 October 2019, we found blue crusts on mid intertidal rocks that directly faced the open sea in a wave-exposed rocky intertidal habitat (42.3555, 10.8670) of the Franco Promontory in western Giglio island, Tuscan Archipelago, Tyrrhenian Sea, Italy. Along this coast, the rocky substrate consists of limestone (Bavestrello et al. 2000). On the same day, we detected a grayish stone-like agglomerate with blue inclusions on a wave-sheltered sandy beach (42.3652, 10.8747) near the Campese resort in Giglio. Using Fourier-transform infrared spectroscopy (FTIR), we later verified these crusts and the agglomerate as ‘plasticrusts’ and ‘pyroplastic’, respectively (see below). While the rocky intertidal habitat is barely accessible to the public, the sandy beach is frequently used for recreational activities including fishing, diving, boating and campfires. From 18 to 20 October 2019, we also surveyed rocky intertidal habitats located in three wave-sheltered bays (Campese Bay East: 42.3657, 10.8747; Campese Bay West: 42.3700, 10.8813; Allume Bay: 42.3528, 10.8802) in Giglio for ‘plasticrust’ occurrence and plastic debris.

### Field and lab work

To measure ‘plasticrust’ density in the rocky intertidal, we haphazardly deployed four quadrats (10 cm × 10 cm) where ‘plasticrusts’ were present. We took pictures of the quadrats to determine ‘plasticrust’ area and percent cover in GIMP 2.10 software. Using a knife, we carefully removed ‘plasticrust’ pieces and determined ‘plasticrust’ piece thickness with a digital microscope (VHX-2000, Keyence, Osaka, Japan). We measured ‘pyroplastic’ size with digital calipers. Using a scalpel, we cut small chips out of the ‘pyroplastic’. We kept ‘plasticrust’ pieces and ‘pyroplastic’ chips for subsequent FTIR analyses.

### Fourier-transform infrared spectroscopy (FTIR)

We identified the ‘plasticrust’ and the ‘pyroplastic’ polymer types through FTIR (Vertex 70, Bruker, Ettlingen, Germany). We conducted our FTIR measurements in attenuated total reflectance (ATR) mode in a wavenumber range between 4000 and 370 cm^−1^ with 8 co-added scans and a spectral resolution of 4 cm^−1^. Then, we compared the obtained spectra with the Bruker spectral library using Opus 7.5 software. A hit quality > 700 hits was set as threshold for polymer identification (Bergmann et al. 2017).

## Results

‘Plasticrust’ (Fig. 1 A-C) density was 3.25 ± 1.65 ‘plasticrusts’/dm^2^ (mean ± SE; n = 4 quadrats). ‘Plasticrust’ area was 0.46 ± 0.08 mm^2^ (n = 13 ‘plasticrusts’). ‘Plasticrust’ cover was 0.02 ± 0.01 % (n = 4 quadrats). ‘plasticrust’ thickness ranged between 0.5 and 0.7 mm. FTIR analyses revealed the ‘plasticrust’ material as polyethylene (PE, Fig. 1 D). We did not detect any other crusts in Campese Bay East, Campese Bay West and Allume Bay.

**Fig. 1.**
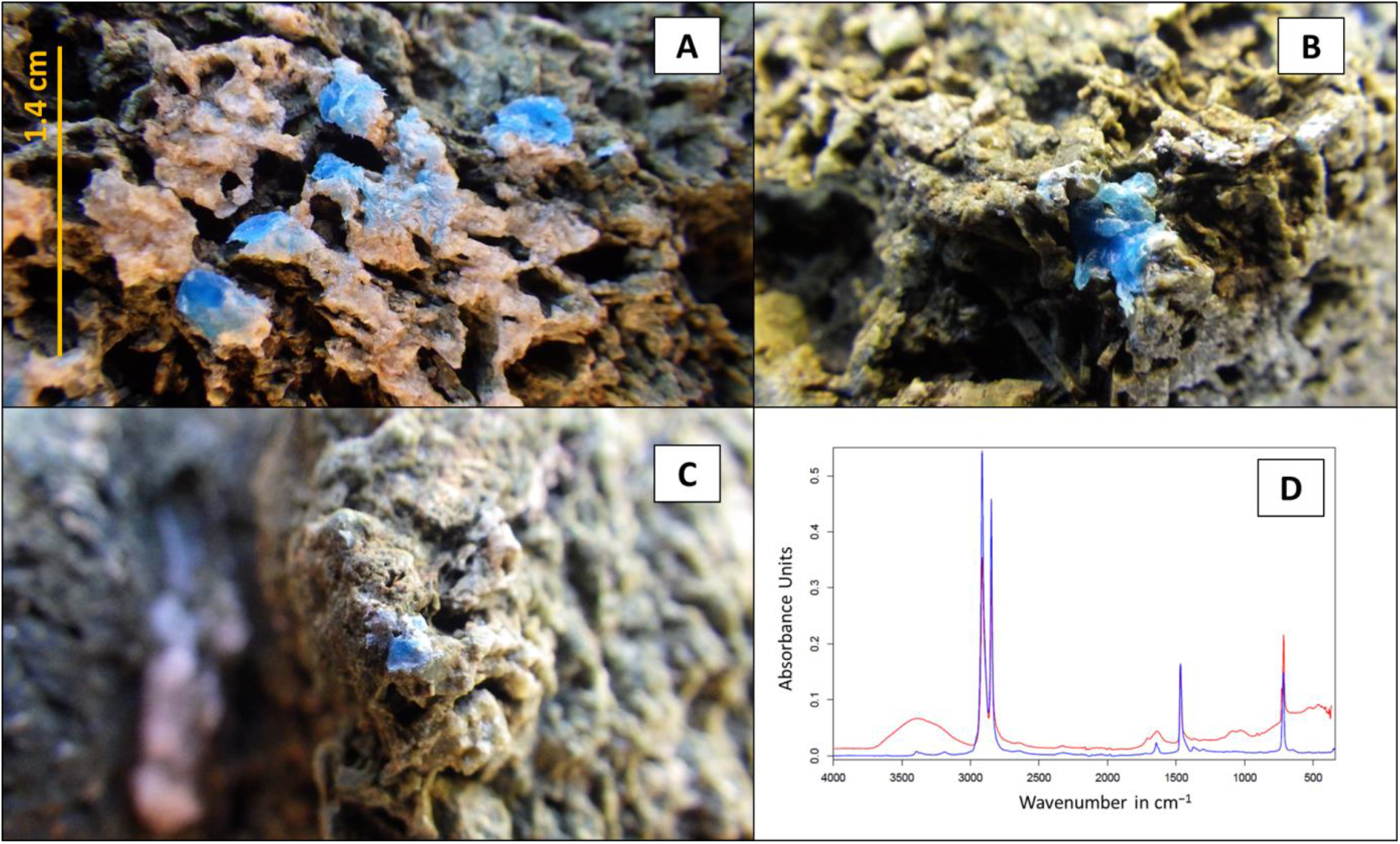
A-C) Blue ‘plasticrusts’ found in Giglio. D) The corresponding FTIR spectrum that revealed polyethylene (PE) as the ‘plasticrust’ material. The red spectrum resulted from the FTIR measurement and the blue spectrum is from the Bruker spectral library.

‘Pyroplastic’ (Fig. 2 A) size was approximately 2 cm × 1.4 cm × 0.5 cm (length × width × height). FTIR analyses showed that the ‘pyroplastic’ material was polyethylene terephthalate (PET, Fig. 2 B). On the sandy beach, we detected burnt charcoal near the ‘pyroplastic’. Finally, we found several blue PET bottles (Fig. 2 C & D) in each surveyed location

**Fig. 2.**
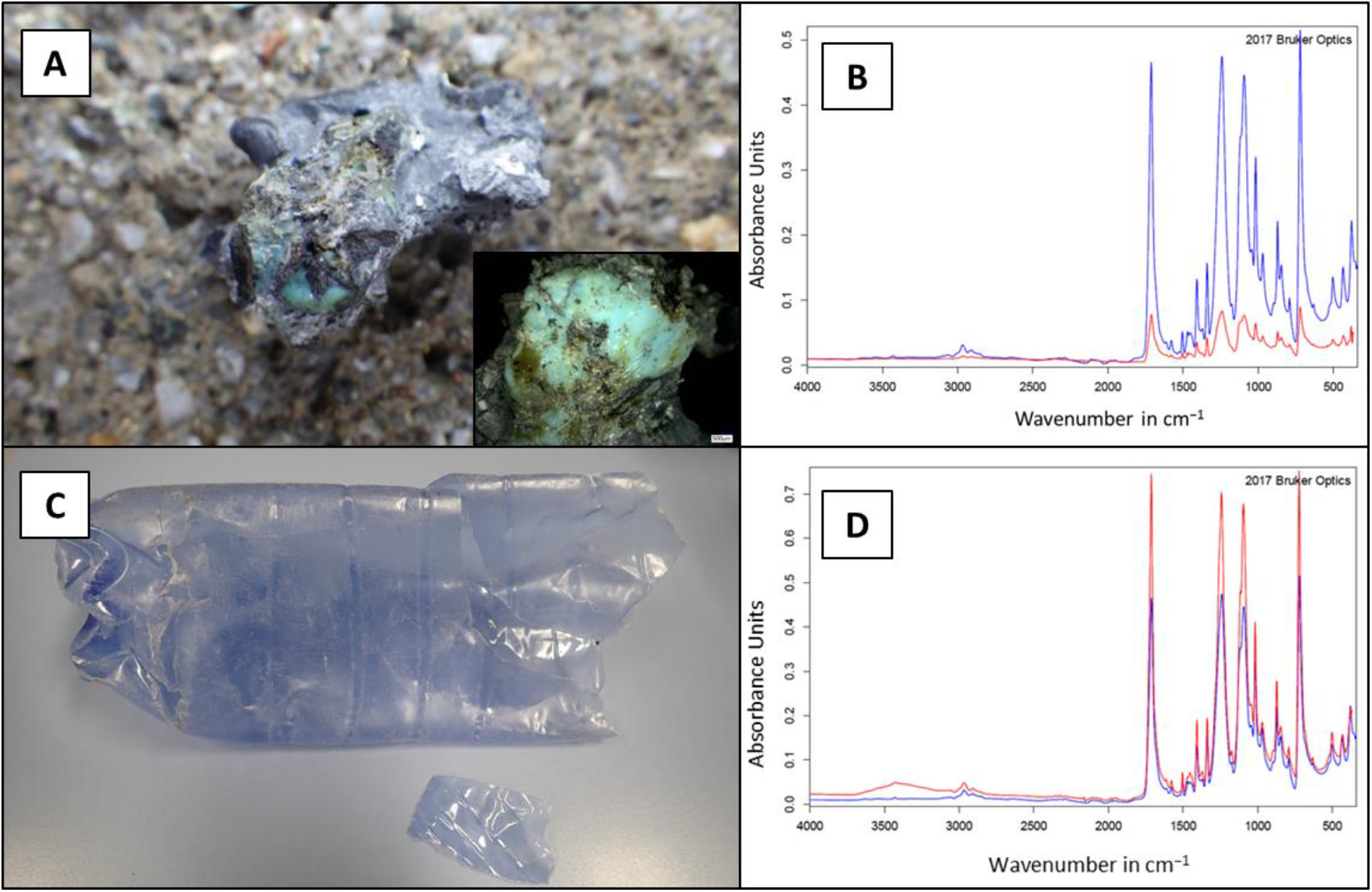
A) The ‘pyroplastic’ with a blue inclusion (see insertion) detected in Giglio. B) The corresponding FTIR spectrum that reveals polyethylene terephthalate (PET) as the ‘pyroplastic’ material. C) A blue beverage bottle found in Giglio. D) The FTIR spectrum that reveals PET as the beverage bottle material. The red spectra resulted from the FTIR measurements and the blue spectra are from the Bruker spectral library.

## Discussion

Our ‘plasticrusts’ results from Giglio island, Tyrrhenian Sea, resembled recent ‘plasticrust’ findings from Madeira island, NE Atlantic Ocean, regarding ‘plasticrust’ color (white and blue), thickness (0.77 ± 0.10 mm, mean ± SE) and material (Polyethylene, PE; Gestoso et al. 2019). Although our plastic debris survey did not detect any blue PE items in Giglio, it is not surprising that the ‘plasticrust’ consisted of PE since this polymer is one of the most frequently used plastics worldwide (Kalogerakis et al. 2017) and, consequentially, PE pollution is common along NE Atlantic (Lots et al. 2017) and Tyrrhenian coastlines (Avio et al. 2017, Baini et al. 2018, Caldwell et al. 2019) including Madeira (Lots et al. 2017) and Giglio (Avio et al. 2017), respectively.

‘Plasticrusts’ are generated by sea waves smashing plastic debris against rugose rocks (Gestoso et al. 2019). Interestingly, ‘plasticrust’ abundance was much higher in Madeira (9.46 ± 1.77 % cover, mean ± SE; Gestoso et al. 2019) than in Giglio (0.02 ± 0.01 % cover) on rocks of similar rugosity (Gestoso et al. 2019, Fig. 1 A-C). Floating plastic debris is common off Madeira (Chambault et al. 2018) and Giglio (Campanale et al. 2019). However, while Madeira is an isolated off-shore island exposed to open Atlantic waves, Giglio is a nearshore island that is relatively wave-sheltered among larger nearby islands (Elba, Corsica and Sardinia) and the Italian mainland. Moreover, wind speed and wave height are typically higher in the NE Atlantic Ocean than in the Tyrrhenian Sea (Gazeau et al. 2004) suggesting that differences in wave exposure between Madeira and Giglio may actually cause the detected difference in ‘plasticrust’ abundance between the two islands. This notion is supported by the fact that we did not detect any other ‘plasticrusts’ during our survey of rocky intertidal habitats in the three wave-sheltered bays (Campese Bay West, Campese Bay East, Allume Bay) in Giglio. Finally, while the Madeira ‘plasticrusts’ were spread across the mid to upper intertidal (Gestoso et al. 2019), the Giglio ‘plasticrusts’ where limited to a relatively narrow strip along the mid intertidal (Fig. 1 A). Since the maximum tidal amplitude in Madeira (ca. 2.6 m; Cacabelos et al. 2019) is much wider than in the Tyrrhenian Sea (ca. 0.45 m; Cutroneo et al. 2017), this observation suggests that tidal amplitude may influence ‘plasticrust’ spread. Thus, we conclude that ‘plasticrust’ abundance may depend on the local levels of nearshore plastic debris, wave exposure and tidal amplitude. We propose that this conclusion could be examined, for instance, by combining environmental modelling and coastal surveys to predict and verify ‘plasticrust’ abundance in order to deepen the understanding for ‘plasticrust’ generation.

Microplastic (particles < 5 mm, Andrady 2011) can enter food webs via consumption by invertebrates, such as snails (Gutow et al. 2016) and crabs (Watts et al. 2014, Brennecke et al. 2015), and can harm the affected organisms (Watts et al. 2015, Brennecke et al. 2015).

For instance, as microplastic is readily ingested by grazing snails (Gutow et al. 2016, Ehlers et al. 2019, Panebianco et al. 2019), it has been suggested that ‘plasticrusts’ may be consumed by co-occurring intertidal snails (*Tectarius striatus*) in Madeira (Gestoso et al. 2019). Along the Tyrrhenian Sea coast, limpets (*Patella aspera*, *Patella rustica*) and topshells (*Phorcus turbinatus*) are common snails grazing on low intertidal and shallow subtidal rocks (Heller 2015, Sousa et al. 2018). In Giglio, we found these snails on rocks ca. 0.5 m below the ‘plasticrusts’ suggesting that they may not contact the ‘plasticrusts’. However, we frequently observed marbled rock crabs (*Pachygrapsus marmoratus*) at the ‘plasticrust’ elevation. These common omnivorous crabs consume algae and invertebrates (Cannicci et al. 2002, Fratini et al. 2013) and may, therefore, represent a potential microplastic pathway into the Tyrrhenian food web.

The ‘pyroplastic’ from Giglio visually resembles those reported from beaches in the SW United Kingdom (UK; Turner et al. 2019) regarding its stone-like appearance, size, grayish color and blue inclusions (Fig. 2 A). However, in contrast to UK ‘pyroplastics’, the Giglio ‘pyroplastic’ did not consist of PE or polypropylene (PP) but of polyethylene terephthalate (PET) which is a typical beverage bottle material (Oberbeckmann et al. 2016). Interestingly, we frequently found blue PET bottles in Giglio and also witnessed burnt charcoal on the beach where we detected the ‘pyroplastic’. These observations suggest that the detected ‘pyroplastic’ may have resulted from a PET bottle that melted during a campfire. Alternatively, the ‘pyroplastic’ may have resulted from a waste incineration fire since we frequently noticed the strong smell of melting plastic from nearby fires on private properties at several occasions during our daytime field surveys.

Upon detecting ‘plasticrusts’ in Madeira, Gestoso et al. (2019) recommended that this new plastic debris type may be included in marine conservation policy and mentioned the need to document ‘plasticrusts’ on other coasts. Similarly, Turner et al. (2019) stressed that information on the distribution of ‘pyroplastics’ is lacking and that ‘pyroplastics’ should be reported from regions worldwide. Our ‘plasticrust’ and ‘pyroplastic’ findings in Giglio show that these novel plastic debris types are not local phenomena. Furthermore, ‘plasticrusts’ and ‘pyroplastics’ may harm invertebrates through microplastic ingestion and toxic substance release, respectively (Gestoso et al. 2019, Turner et al. 2019). Therefore, we recommend that risk assessment studies should document the distribution and abundance of these novel plastic debris types and examine the ecotoxicity of these potentially hazardous materials in order to decide upon ‘plasticrust’ and ‘pyroplastics’ inclusion in marine conservation policy.

## Acknowledgments

We thank Jochen HE Koop (Department of Animal Ecology, Federal Institute of Hydrology, Koblenz, Germany) for letting us use the Fourier-transform infrared spectrometer (FTIR) for polymer type identification. This research did not receive any specific grant from funding agencies in the public, commercial, or not-for-profit sectors.

